# Bacteriophage-mediated lysis supports robust growth of amino acid auxotrophs

**DOI:** 10.1101/2023.02.28.530524

**Authors:** Gordon J. Pherribo, Michiko E. Taga

**Affiliations:** Department of Plant & Microbial Biology, University of California, Berkeley, Berkeley, CA 94720 U.S.A.

## Abstract

The majority of microbes are auxotrophs – organisms unable to synthesize one or more metabolites required for their growth. Auxotrophy is thought to confer an evolutionary advantage, yet auxotrophs must rely on other organisms that produce the metabolites they require. The mechanisms of metabolite provisioning by “producers” remain unknown. In particular, it is unclear how metabolites such as amino acids and cofactors, which are found inside the cell, are released by producers to become available to auxotrophs. Here, we explore metabolite secretion and cell lysis as two distinct possible mechanisms that result in release of intracellular metabolites from producer cells. We measured the extent to which secretion or lysis of *Escherichia coli* and *Bacteroides thetaiotaomicron* amino acid producers can support the growth of engineered *Escherichia coli* amino acid auxotrophs. We found that cell-free supernatants and mechanically lysed cells provide minimal levels of amino acids to auxotrophs. In contrast, bacteriophage lysates of the same producer bacteria can support as many as 47 auxotroph cells per lysed producer cell. Each phage lysate released distinct levels of different amino acids, suggesting that in a microbial community the collective lysis of many different hosts by multiple phages could contribute to the availability of an array of intracellular metabolites for use by auxotrophs. Based on these results, we speculate that viral lysis could be a dominant mechanism of provisioning of intracellular metabolites that shapes microbial community structure.

## Main Text

Microbial communities are ubiquitous on earth, contributing to global nutrient cycling, human health and agricultural productivity. Nutritional interdependence is a universal feature of microbial communities [1]. One form of nutritional interaction occurs between auxotrophs – organisms unable to produce a nutrient required for growth – and prototrophs or “producers” – organisms capable of synthesizing a nutrient required by others. Auxotrophy contributes to microbial community stability [2, 3], and genomic analysis suggests that the majority of bacteria are auxotrophic for one or more amino acids or cofactors [4, 5], likely because auxotrophy often confers a growth advantage when the required nutrient is available [5]. The prevalence of auxotrophy suggests that a consistent source of nutrients must be available to auxotrophs over evolutionary timescales, yet it is unknown how these nutrients become available. Here, we investigate how metabolites that function intracellularly, such as amino acids and cofactors, become accessible to auxotrophs. We previously proposed that nutrient release resulting from secondary effects of evolved biological processes, such as secretion of excess metabolites or cell lysis, are possible mechanisms of providing nutrients to auxotrophs [6]. Here, we experimentally test both mechanisms using two phylogenetically distinct producer species and engineered *Escherichia coli* amino acid auxotrophs.

Amino acids and dipeptides have been found in exometabolomes of cultured microbes and soil biocrusts, suggesting they are available to auxotrophs in these environments [7, 8]. Furthermore, certain pairs of amino acid auxotrophs can grow in coculture in the absence of amino acid supplementation, suggesting that bacteria may shed amino acids during growth [4, 9, 10]. To test whether amino acid secretion or leakage can be sufficient to support the growth of amino acid auxotrophs, we measured the extent to which cell-free supernatants from *E. coli* and *Bacteroides thetaiotaomicron,* both producers of all 20 proteogenic amino acids, could support the growth of up to 11 different *E. coli* amino acid auxotrophs in the absence of amino acid supplementation. We found that supernatants from both *E. coli* and *B. thetaiotaomicron* minimally supported most of the tested amino acid auxotrophs used in this study, with each producer cell supporting the growth of less than 0.2 auxotroph cells in all except one case (Fig. 1A and 1B). The exception was the tryptophan (Trp) auxotroph, in which each *E. coli* producer cell was estimated to support about one Trp auxotroph. Amino acid provisioning was only slightly influenced by growth phase, with modest levels of auxotroph growth observed for producers harvested in stationary (Fig. 1) and exponential phase (Fig. S1). Together, these results suggest that secretion or leakage may not be dominant forms of amino acid provisioning.

**Figure 1.**
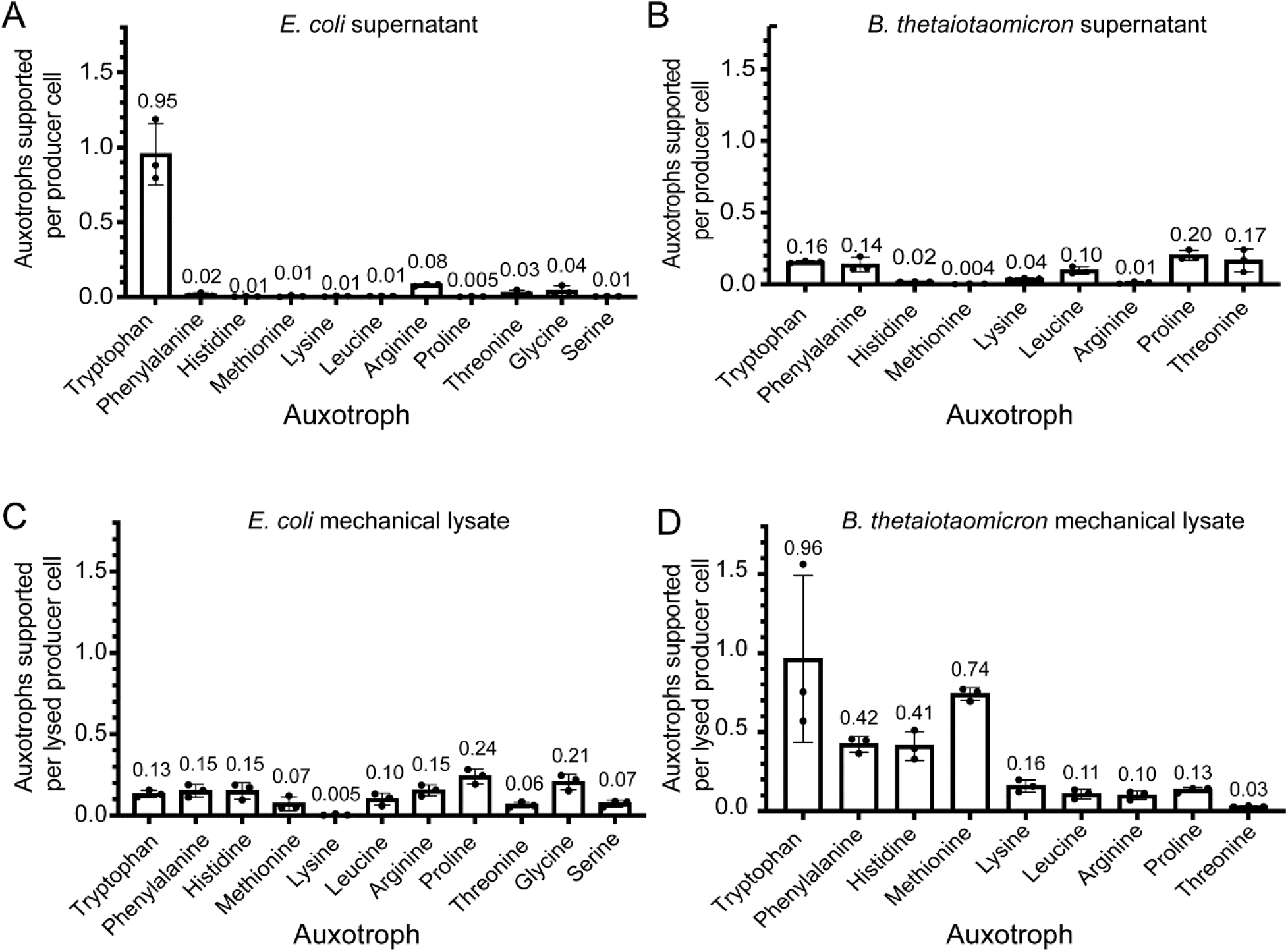
Cell-free supernatants and mechanical lysates minimally support growth of amino acid auxotrophs. The number of auxotroph cells supported per producer cell was calculated for auxotrophs grown in defined medium containing (A, B) cell-free supernatants and (C, D) French pressed lysates of (A, C) *E. coli* and (B, D) *B. thetaiotaomicron.* The auxotrophs are ordered from left to right on the x-axis based on the estimated biosynthetic cost of producing one molecule of the amino acid [4]. The genotype of each auxotroph is listed in Supplemental Table 1. In all except one case (Met), a single gene in the biosynthetic pathway is deleted [4].

We next explored the possibility that cell lysis can release amino acids at sufficient levels to support auxotrophs. Microbial cell lysis occurs naturally by a variety of mechanisms including exposure to toxins or developmental genetic programs, such as sporulation in *Bacillus subtilis* and fruiting body formation in *Myxcococcus xanthus* [11–13]. A common trigger of microbial lysis is viral infection, as viruses are abundant and ubiquitous across many environments [14]. Environmentally, viruses are major drivers of microbial turnover and elemental cycling in marine environments, a process known as the viral shunt [15, 16]. Further, in a laboratory coculture, addition of a lytic phage can lead to increased growth yield of a partner strain [17, 18]. Thus, there is evidence for cell lysis as a mechanism of nutrient provisioning in both ecological systems and laboratory coculture.

To determine whether the levels of intracellular amino acids liberated via lysis of a producer could be sufficient to support the growth of amino acid auxotrophs, we tested the ability of the 11 amino acid auxotrophs to grow in media supplemented with mechanically lysed *E. coli* or *B. thetaiotaomicron* cells. We observed that lysates of both producers contain a higher level of bioavailable amino acids than their respective supernatants, though less than one auxotroph cell was supported by each lysed producer cell (Fig. 1C and D). Comparison of lysates of *E. coli* and *B. thetaiotaomicron* suggests that these bacteria contain different levels of several amino acids, which likely reflects their distinct metabolic characteristics. Interestingly, the strains that were most supported by mechanical lysates of *B. thetaiotaomicron* were those auxotrophic for the more biosynthetically costly amino acids (Fig. 1D). Yet, with less than one auxotroph cell supported for each lysed producer cell, the ratio of producer to auxotroph cells in a community would need to be high.

Given that viral infection reprograms host metabolism, different levels of amino acids become available via phage lysis as compared to mechanical lysis [19, 20]. We therefore measured how lysis of producers following phage infection shapes amino acid bioavailability. *E. coli T4rI* phage lysates supported the growth of all 11 auxotrophs at levels higher than supernatants and mechanically lysed *E. coli,* with nine of the 11 auxotrophs at a level exceeding one auxotroph supported by one producer (Fig. 2A). Remarkably, five auxotrophs were supported at levels ranging from 11 to 47 auxotrophs per producer, suggesting that a low ratio of producer to auxotroph cells could coexist (Fig. 2A).

**Figure 2.**
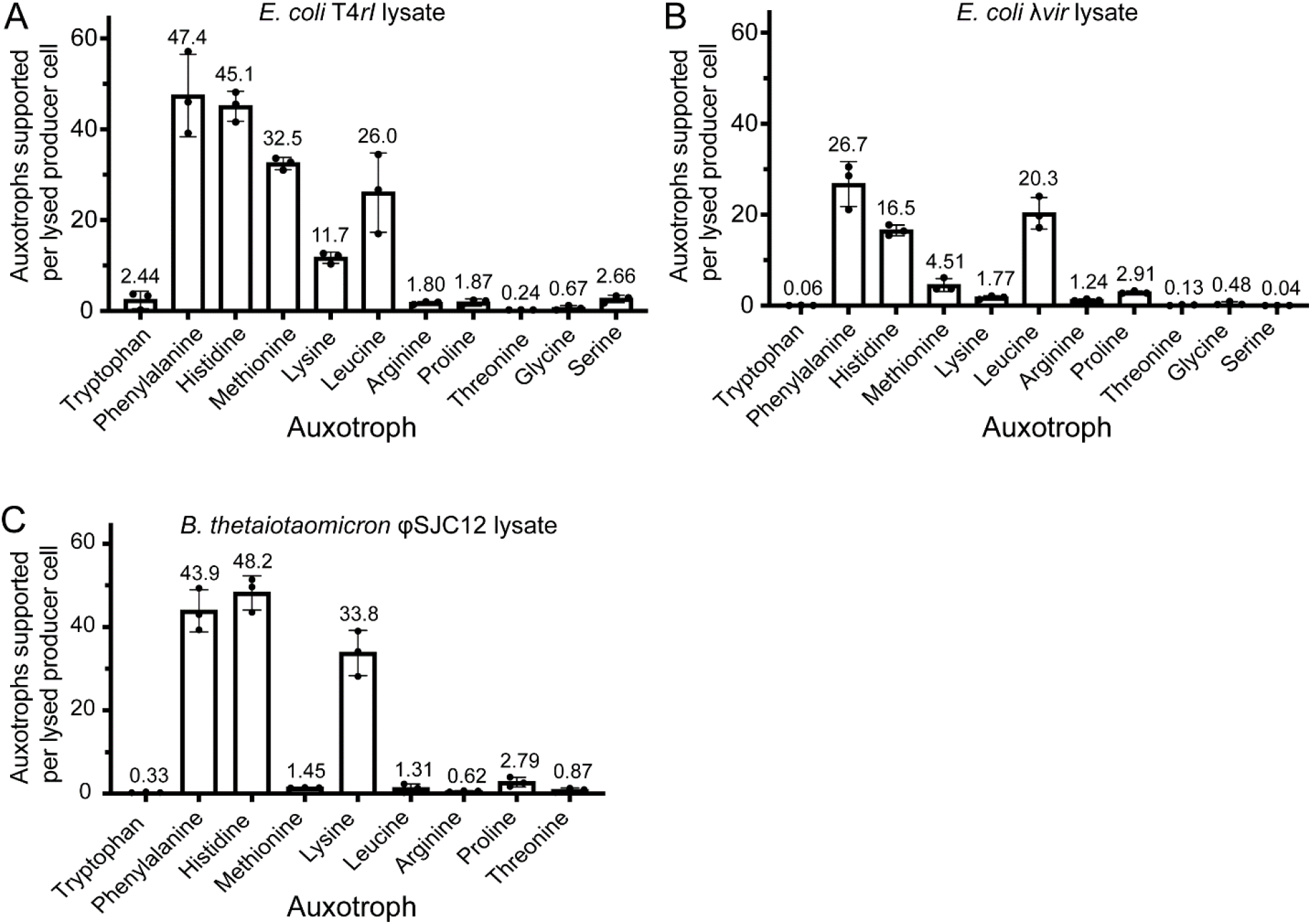
Phage lysates robustly support growth of certain amino acid auxotrophs. The number of auxotroph cells supported per producer cell was calculated for auxotrophs grown in defined medium containing (A) *E. coli T4rI* phage lysate, (B) *E. coli λvir* lysate and (C) *B. thetaiotaomicron* φSJC12 lysate. The amino acids are ordered as described in Fig. 1.

Similar to the *T4rI* phage lysates, *E. coli λvir* lysates contained high concentrations of certain amino acids, supporting as many as 27 auxotrophs per producer cell (Fig. 2B). Upon observing that the two *E. coli* phage lysates showed differences in the levels of bioavailable amino acids, we compared these results with nine auxotrophs treated with lysates of *B. thetaiotaomicron* infected with the lytic phage ΦSJC12 and found they supported auxotroph growth to a higher degree than *B. thetaiotaomicron* supernatants or mechanical lysates with, in three cases, over 30 auxotroph cells supported per producer cell (Fig. 2C). These results demonstrate that (1) phage lysis contributes a higher level of most amino acids compared to mechanical lysis or secretion, (2) different phage infections and host cells vary in the levels of bioavailable amino acids released upon lysis, and (3) in general, biosynthetically costly amino acids are released at higher levels than less costly amino acids. Though these results are limited to two producer bacteria and three phages in artificial conditions, they suggest that infection of diverse bacteria with different phages could support a variety of auxotrophs in microbial ecosystems. We postulate that phage lysis may be a dominant mechanism of intracellular nutrient release, potentially shaping the nutritional environment and contributing to the evolution of auxotrophy.

## Acknowledgements

We thank Richard Calendar (in memoriam) and Drew Hryckowian for providing phage stocks. We are grateful to members of the Taga lab for helpful discussions, the UC Berkeley Department of Statistics consulting service for advice on statistical analysis, and Janani Hariharan, Alexa Nicolas, and Zachary Hallberg for critical reading of the manuscript. This work was funded by an NSF doctoral fellowship to G.J.P. and NIH grant R35GM139633 to M.E.T.

