## Supplemental materials for "Bacteriophage-mediated lysis supports robust growth of amino acid auxotrophs"

### Materials & Methods

#### Bacterial culturing and media

*E. coli* strains were inoculated from single colonies on LB plates into 2 mL H1 glycerol minimal medium [21] and grown with aeration at 200 rpm overnight at 37 °C prior to all experiments. *E. coli* auxotrophs were supplemented with their required amino acid for all pre-culturing (methionine: 0.2 mg/mL, arginine: 0.2 mg/mL, proline: 0.5 mg/mL, glycine: 0.32 mg/mL, histidine: 0.625 mg/mL, phenylalanine: 0.2 mg/mL, lysine: 0.3 mg/mL, threonine: 0.2 mg/mL, leucine: 0.03 mg/mL, serine: 1.5 mg/mL, glycine: 0.625 mg/mL, tryptophan: 0.5 mg/mL).

*B. thetaiotaomicron* strains were inoculated from single colonies on Bacteroides Phage Recovery Medium (BPRM) plates [22] into 5 mL BPRM or defined minimal medium at 37 °C in an anaerobic chamber (Coy Labs) containing 10% $H_2$, 10% $CO_2$, and 80% $N_2$. The defined minimal medium was prepared as described [23], except that sucrose was substituted for glucose and 10 nM vitamin $B_{12}$ was substituted for methionine.

#### Preparation of supernatants and mechanical lysates

Cultures of *E. coli* MG1655 in H1 glycerol minimal medium were grown to saturation and diluted 1:100 in fresh H1 glycerol minimal medium. Once the cultures reached an optical density ($O.D._{600}$) of 0.3 to 0.4, cells were harvested by centrifugation at 4,427 rcf for 10 minutes. The supernatant was passed through a 0.2 um filter and stored at -80 °C. The cell pellet was washed three times with H1 glycerol minimal medium, resuspended in fresh medium, and cells were lysed using a French Press (Thermo Fisher Scientific). Uninoculated medium used as a control was also subjected to French press. French pressed samples were centrifuged for 15 minutes at 6000 rcp to remove cell debris, and the supernatant was sterilized by passing through a 0.2 um filter and stored at -80 °C. Mechanical lysates of *B. thetaiotaomicron* were generated similarly, except this strain was cultured anaerobically in defined minimal medium. Each lysate was generated in triplicate.

The number of cells lysed by French press was calculated by removing aliquots of cultures before and after French press treatment, serially diluting in 96 well plates, and spot plating each dilution. The number of cells lysed was calculated as the difference in colony forming units (CFU)/ml before and after lysis. This measurement was performed in triplicate.

#### One-Step Phage Growth Curve

Phages used in this study are listed in Supplemental Table 2. The one-step phage growth curve was used to determine the average phage burst size and the amount of time required to generate ϕSJC12 lysates for the auxotroph assay [24]. One-step growth curves were performed in triplicate as in [24]. Briefly, *B. thetaiotaomicron* was grown in defined minimal medium at 37 °C in the anaerobic chamber (10% $H_2$, 10% $CO_2$, and 80% $N_2$) to a cell density of approximately $10^7$ CFU/mL. The ϕSJC12 phage lysate was added to obtain a multiplicity of infection (MOI) of 0.01 ($10^5$ PFU/mL). The bacteria-phage mix was incubated for 10 min at room temperature to allow for phage adsorption and then transferred to 37 °C. At each time point, 0.4 ml was removed. Half of the sample was added to a few drops of chloroform, vortexed for 10 seconds, and placed at room temperature until the chloroform settled. The remaining 0.2 ml was stored on ice and processed during the experiment. These samples were serially diluted up to $10^{-8}$, and spot plated onto BPRM top agar overlays (0.35% w/v) prepared with *B. thetaioaomicron* host cells to measure infective centers and free phage, and plated onto BPRM agar (1.5% w/v) to measure cell survival following phage addition. Three biological replicates were measured for each growth curve. At the end of the experiment, chloroform-treated samples were serially diluted up to $10^{-8}$, spot plated onto BPRM top agar overlays, and grown overnight in an anaerobic chamber. One-step growth curves for T4*rI* and λ*vir* were performed similarly, except that *E. coli* cultures were grown aerobically in H1 glycerol minimal medium at 37°C. For PFU calculations, samples were spot plated on LB top agar overlays prepared with *E. coli* host cells (0.7% w/v) and CFU measurements were spot plated on LB agar (1.5% w/v).

**Generating Phage Lysates for Auxotroph Assay**

To generate *E. coli* phage lysates for the auxotroph assay, saturated cultures of *E. coli* MG1655 were diluted 1:100 in fresh H1 glycerol minimal medium. Once cells reached an $O.D._{600}$ of approximately 0.3, T4*rI* phage was added at an MOI of 3 for 100 minutes. Each phage lysate treatment had a parallel mock treatment that served as a control, prepared by passing phage stocks twice through 100 kDa centrifugal filters (Amicon). After 100 minutes, the phage-treated culture and the mock treatment were collected by centrifugation at 4,427 rcf for 10 minutes, and bacterial cells were removed by passing through a 0.2 um filter. The number of new phage particles generated during incubation was calculated as the difference between the titer measured at the beginning and end of the incubation period. This value was divided by the burst sizes calculated from the one-step growth curves to estimate the number of cells lysed by the phage. The samples were further processed by passing through a 100 kDa centrifugal filter to remove phage particles. Samples were stored at -80 °C. Phage lysates for λ*vir* were prepared similarly except the incubation time was 150 minutes. Phage lysates from *B. thetaiotaomicron* were similarly processed, except the bacteria were grown anaerobically at 37°C in defined minimal medium, and *B. thetaiotaomicron* was incubated with φSJC12 for 4 hours. All lysates were generated in triplicate.

***E. coli* Auxotroph Assay**

*E. coli* amino acid auxotrophs were pre-cultured in biological triplicate in H1 glycerol minimal medium at 37°C with their required amino acid for 18-24 hours until saturation, and subsequently washed three times in H1 glycerol minimal medium to remove amino acids supplemented during the pre-culturing step. The cultures were then diluted to an $O.D._{600}$ of 0.02 in H1 glycerol minimal medium, and 100 μl of culture was dispensed into wells of a 96-well plate (Corning ®). One hundred microliters of phage lysate was then added to each well, bringing the final $O.D._{600}$ to 0.01 and the final glycerol concentration to 0.2%. The plates were shaken at 1,200 rpm for 24 hours at 37°C in a Southwest Science heated plate shaker. Absorbance measurements were measured using a multi-well plate reader (Tecan Spark). Colony forming units (CFU) were calculated after 24 hours of growth by spot plating 10-fold serial dilutions onto LB agar plates.

**Phage Lysate Preparation and Titering**

*E. coli* MG1655 was used as the host for the propagation of T4*rI* (Carolina Biological Supply Company). High-titer stocks of *E. coli* phages were generated by growing cultures at 37 °C with aeration in either LB or H1 glycerol minimal medium to an $O.D._{600}$ of 0.2-0.3, and adding phage to an MOI of 0.01. After about 7 hours, chloroform was added to lyse remaining cells. After the chloroform settled, the sample was centrifuged at 6,026 rcf and the lysate was passed through a 0.2 um filter to remove bacterial cells. Phage stocks were titered by spot plating onto soft agar overlays. High titers for λ*vir* were generated similarly.

*B. thetaiotaomicron* was used as the host for the propagation of φSJC12 phage. φSJC12 was a gift from Andrew Hryckowian and Bryan Merrill. High-titer stocks of φSJC12 were generated by a soft agar overlay method similar to [25]. Briefly, 0.5 ml of saturated *B. thetaiotaomicron* culture was combined with 1-10 µl of high-titer φSJC12 stock for 20 minutes to allow for phage adsorption. Then 4.5 mL of molten BPRM top agar (0.35%) prepared with *B. thetaiotaomicon* host cells added was poured onto BPRM agar (1.5 %) and incubated anaerobically overnight at 37°C. Top agar overlays that showed a "lacy" pattern (confluent lysis) were flooded with sterile phage buffer (an autoclaved solution of 5 mL of 1 M Tris pH 7.5, 2 g NaCl, 5 mL of 1 M $MgSO_4$, in 500 mL of $ddH_2O$) and incubated at room temperature for at least 2 hours to suspend the phage. The phage buffer was removed, passed through a 0.2 µm filter and stored at 4 °C for future experiments. For phage lysates used to generate lysate treatments for the auxotroph assays, defined minimal medium top agar was used instead of BPRM top agar.

**Calculating Amino Acid Auxotrophs per Lysed Cell**

To calculate the number of auxotrophs supported per lysed cell, the difference in CFU/ml between the treatment (addition of supernatant, French pressed lysate, or phage lysate) and its control (no addition control, French press control, or mock phage) was divided by the calculated number of lysed cells.

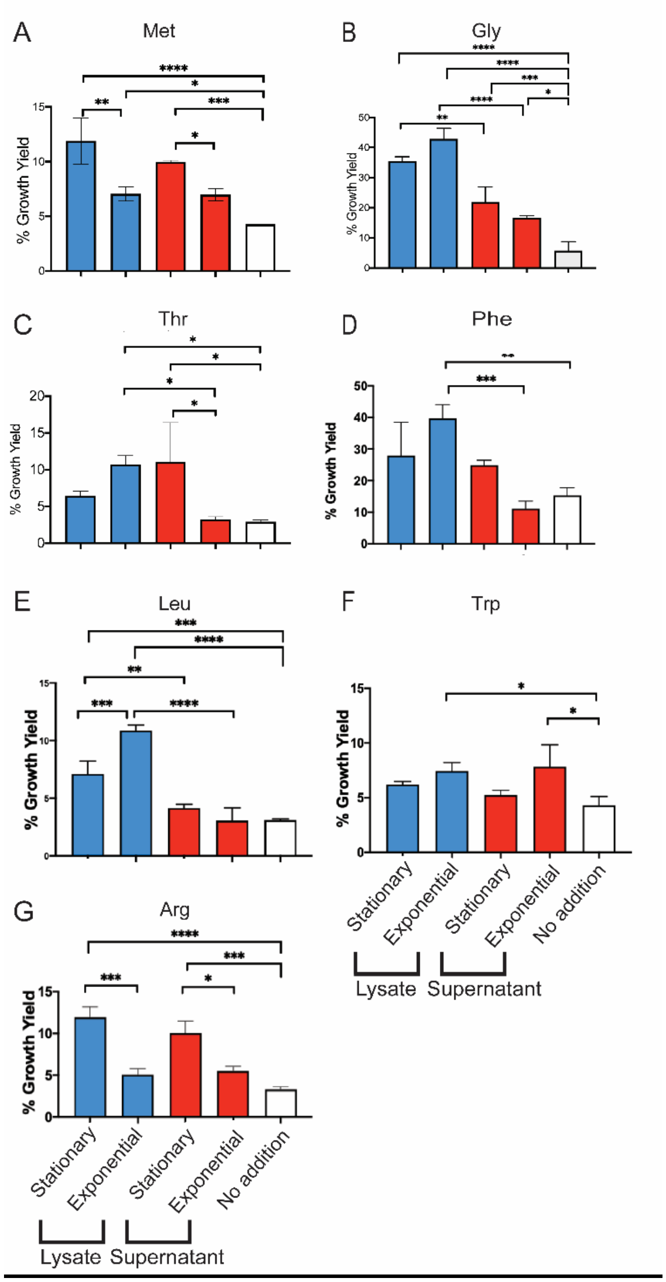


Supplemental Figure 1. The growth yield of each amino acid auxotroph was measured after treatment with lysates and supernatants from either stationary or exponential phase *E. coli* cultures. Growth yield was calculated as the $O.D._{600}$ of cultures grown with the indicated supplements as a percentage of the $O.D._{600}$ of cultures grown with a saturating concentration of the required amino acid. Significance was determined by a one-way ANOVA with Tukey's multiple comparison test ($p<0.0001$ = ****; $p<0.0002$= ***, $p<0.0021$=**, $p<0.0332$= *)

**Supplemental Table 1. List of bacterial strains used in this study.**

| Species | Strain | Auxotrophy | Genotype | Reference |
|---|---|---|---|---|
| *E. coli* | MG1655 | Wild type | | This study |
| *B. thetaiotaomicron* | VPI5482 | Wild type | | This study |
| *E. coli* | JW2786-1 | Arginine | BW25113 *ΔargA::kan* | [21] |
| *E. coli* | GP253 | Methionine | MG1655 *ΔmetA, ΔmetB, ΔmetE, ΔmetH, ΔmetC::kan* | This study |
| *E. coli* | JW2535-1 | Glycine | BW25113 *ΔglyA::kan* | (Baba et al, 2006) |
| *E. coli* | JW2004-1 | Histidine | BW25113 *ΔhisB::kan* | (Baba et al, 2006) |
| *E. coli* | JW5807-2 | Leucine | BW25113 *ΔleuB::kan* | (Baba et al, 2006) |
| *E. coli* | JW2806-2 | Lysine | BW25113 *ΔlysA::kan* | (Baba et al, 2006) |
| *E. coli* | JW2580-1 | Phenylalanine | BW25113 *ΔpheA::kan* | (Baba et al, 2006) |
| *E. coli* | JW0233-2 | Proline | BW25113 *ΔproA::kan* | (Baba et al, 2006) |
| *E. coli* | JW2880-1 | Serine | BW25113 *ΔserA::kan* | (Baba et al, 2006) |
| *E. coli* | JW0003-2 | Threonine | BW25113 *ΔthrC::kan* | (Baba et al, 2006) |
| *E. coli* | JW1254-2 | Tryptophan | BW25113 *ΔtrpC::kan* | (Baba et al, 2006) |

**Supplemental Table 2. List of phages used in this study.**

| Phage | Host | Source |
|---|---|---|
| T4*rI* | *E. coli* | Carolina Biological Supply Company |
| λ*vir* | *E. coli* | This study |
| φSJC12 | *B. thetaiotaomicron* | [22] |
